# RAPID: an interactive R/Shiny platform for end-to-end 16S rRNA and ITS amplicon sequence analysis using DADA2

**DOI:** 10.64898/2026.05.05.723040

**Authors:** Beant Kapoor, Melissa A. Cregger, Priya Ranjan

**Author notes:** This manuscript has been authored by UT-Battelle, LLC under Contract No. DE-AC05-00OR22725 with the U.S. Department of Energy. The United States Government retains and the publisher, by accepting the article for publication, acknowledges that the United States Government retains a non-exclusive, paid-up, irrevocable, world-wide license to publish or reproduce the published form of this manuscript, or allow others to do so, for United States Government purposes. The Department of Energy will provide public access to these results of federally sponsored research in accordance with the DOE Public Access Plan (http://energy.gov/downloads/doe-public-access-plan).

## Abstract

**Motivation:** Amplicon sequencing of 16S rRNA and internal transcribed spacer (ITS) gene regions is the most widely used approach for characterizing bacterial and fungal communities, respectively. The DADA2 pipeline has become a standard for inferring amplicon sequence variants (ASVs), offering single-nucleotide resolution over traditional OTU clustering. However, executing the full DADA2 workflow requires proficiency in R programming and manual coordination of multiple sequential steps, presenting a substantial barrier for researchers in clinical, environmental, and agricultural sciences who lack computational training.

**Results:** We present RAPID (R-based Amplicon Pipeline for Interactive DADA2), a pair of R/Shiny applications providing complete graphical user interfaces for 16S rRNA and ITS amplicon sequence analysis. The 16S application implements a 10-step guided workflow from raw paired-end FASTQ files through quality filtering, error learning, dereplication, paired-read merging, chimera removal, taxonomy assignment (SILVA), phyloseq construction with data transformation (rarefaction, relative abundance, or CLR), interactive visualization (rarefaction curves, alpha diversity, NMDS, PCoA, taxonomic abundance), PERMANOVA, and ANCOM-BC2 differential abundance analysis. The ITS application extends this to an 11-step workflow, adding an automated primer removal step using cutadapt with support for multiple primers and length-variable amplicons, and uses the UNITE database for fungal taxonomy. Both applications feature asynchronous background processing, session persistence, real-time progress monitoring, publication-ready figure export, and comprehensive result downloads.

**Availability:** RAPID is freely available at https://github.com/beantkapoor786/RAPID. Both applications can be installed locally on any system with R (version 4.0 or higher) and run as local web applications accessible through a standard browser.

## 1. Introduction

High-throughput amplicon sequencing of marker genes has become the cornerstone of microbial ecology (Hugerth & Andersson, 2017; Zhou et al., 2015). For bacteria and archaea, the 16S ribosomal RNA gene is the standard target, while the internal transcribed spacer (ITS) region is used for fungal community profiling (Schoch et al., 2012). The field has seen a paradigm shift from operational taxonomic unit (OTU) clustering at 97% sequence identity toward exact amplicon sequence variants (ASVs), which provide single-nucleotide resolution and improved reproducibility across studies (Callahan et al., 2017).

The DADA2 algorithm has emerged as one of the most widely adopted tools for ASV inference from Illumina amplicon data (Callahan et al., 2016). By modeling sequencing run-specific error rates, DADA2 distinguishes true biological sequences from artifacts without arbitrary similarity thresholds. DADA2 is available as an R/Bioconductor package and can also be accessed through the QIIME 2 platform (Bolyen et al., 2019). Importantly, DADA2 supports both 16S and ITS workflows, though the ITS pipeline requires additional steps for primer removal due to the highly variable length of ITS amplicons (200-600 bp), which precludes fixed-length truncation and necessitates external tools such as cutadapt for primer removal (Martin, 2011).

Despite the methodological advances offered by DADA2, a significant accessibility gap persists. Executing the complete pipeline requires users to write R scripts, manage package dependencies across CRAN and Bioconductor, configure platform-specific parallelization settings, manually inspect quality profiles, download and manage taxonomy reference databases, and construct phyloseq objects for downstream ecological analysis (McMurdie & Holmes, 2013). For ITS workflows, the additional complexity of primer removal with cutadapt, handling variable-length amplicons, and using the UNITE database further raises the barrier to entry (Nilsson et al., 2019).

Several tools have addressed parts of this challenge. QIIME 2 provides a comprehensive command-line framework but requires terminal proficiency. Shiny-phyloseq offers interactive visualization of processed microbiome data but does not handle upstream sequence processing (McMurdie & Holmes, 2015). MOCHI provides a GUI for amplicon analysis but uses a different processing engine (Zheng et al., 2022). Galaxy-based workflows offer web-based interfaces but require server infrastructure (Batut et al., 2018). None of these tools provide a self-contained, locally installable graphical interface that covers the entire workflow from raw FASTQ files to statistical testing and publication-ready visualizations using the DADA2 algorithm, for both bacterial (16S) and fungal (ITS) marker genes.

Here, we present RAPID, a pair of R/Shiny applications that wrap the complete DADA2 pipeline and downstream analysis into guided, point-and-click graphical user interfaces for both 16S rRNA and ITS amplicon sequencing data. The applications share a common architectural framework but are tailored to the specific requirements of each marker gene, including ITS-specific primer removal and the absence of fixed-length truncation.

## 2. Results

### 2.1 Shared application architecture

Both RAPID applications were developed using the R Shiny framework and share a common architecture featuring (Jia et al., 2022): (i) a step-by-step guided workflow with contextual controls and result summaries at each stage; (ii) asynchronous computation via callr background subprocesses, enabling multicore DADA2 parallelism while maintaining full interface responsiveness; (iii) automatic session persistence to the data directory, allowing users to close and resume analyses across sessions; (iv) a modern dark-themed interface with real-time progress indicators; and (v) publication-ready figure export at 300 DPI. All computationally intensive steps (quality filtering, error learning, dereplication, chimera removal, taxonomy assignment, and ANCOM-BC2) are offloaded to background R subprocesses, resolving the fundamental incompatibility between DADA2’s fork-based multiprocessing and Shiny’s reactive framework.

### 2.2 The 16S rRNA application (10 steps)

The 16S application implements a 10-step workflow: (1) Setup and file detection with auto-identification of forward/reverse read patterns and configurable sample name extraction (**Figure 1**); (2) quality profile visualization with downloadable plots; (3) quality filtering and trimming with all DADA2 parameters exposed (truncLen, maxEE, truncQ, maxN, rm.phix) and post-filter quality verification (**Supplementary Figure 1**); (4) error rate learning and sequence dereplication (**Supplementary Figure 2**); (5) paired-read merging and chimera removal with a read-tracking table (**Supplementary Figures 3 and 4**); (6) taxonomy assignment using the SILVA v138.1 reference database with support for both the naive Bayesian classifier and IdTaxa (DECIPHER) (Murali et al., 2018; Quast et al., 2013; Wang et al., 2007) (**Supplementary Figure 5**); (7) phyloseq object construction with metadata upload, variable type selection, and data transformation (rarefaction to even depth with configurable threshold, relative abundance normalization, or centered log-ratio transformation via the microbiome package) (**Supplementary Figure 6**); (8) interactive visualization including rarefaction curves, alpha diversity boxplots (Observed, Shannon, Simpson), NMDS and PCoA ordination with 95% confidence ellipses and variance-explained axes (**Figure 2**); (9) PERMANOVA with user-specified formula, betadisper homogeneity of dispersion testing, and pairwise comparisons with Bonferroni correction (**Supplementary Figure 7**); and (10) ANCOM-BC2 differential abundance analysis (**Supplementary Figure 7**).

**Figure 1.**
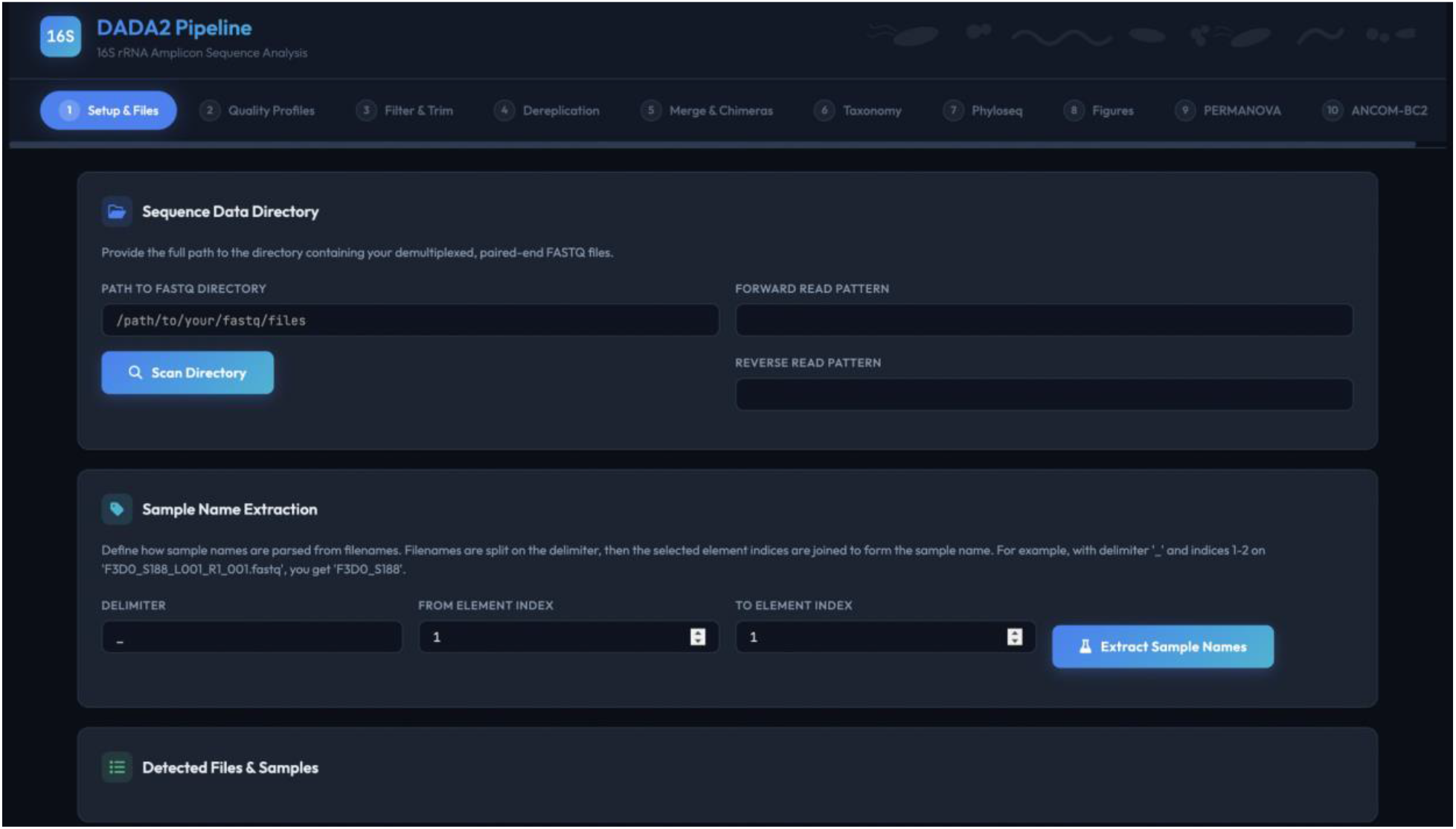
Screenshot of the setup and file detection step.

**Figure 2.**
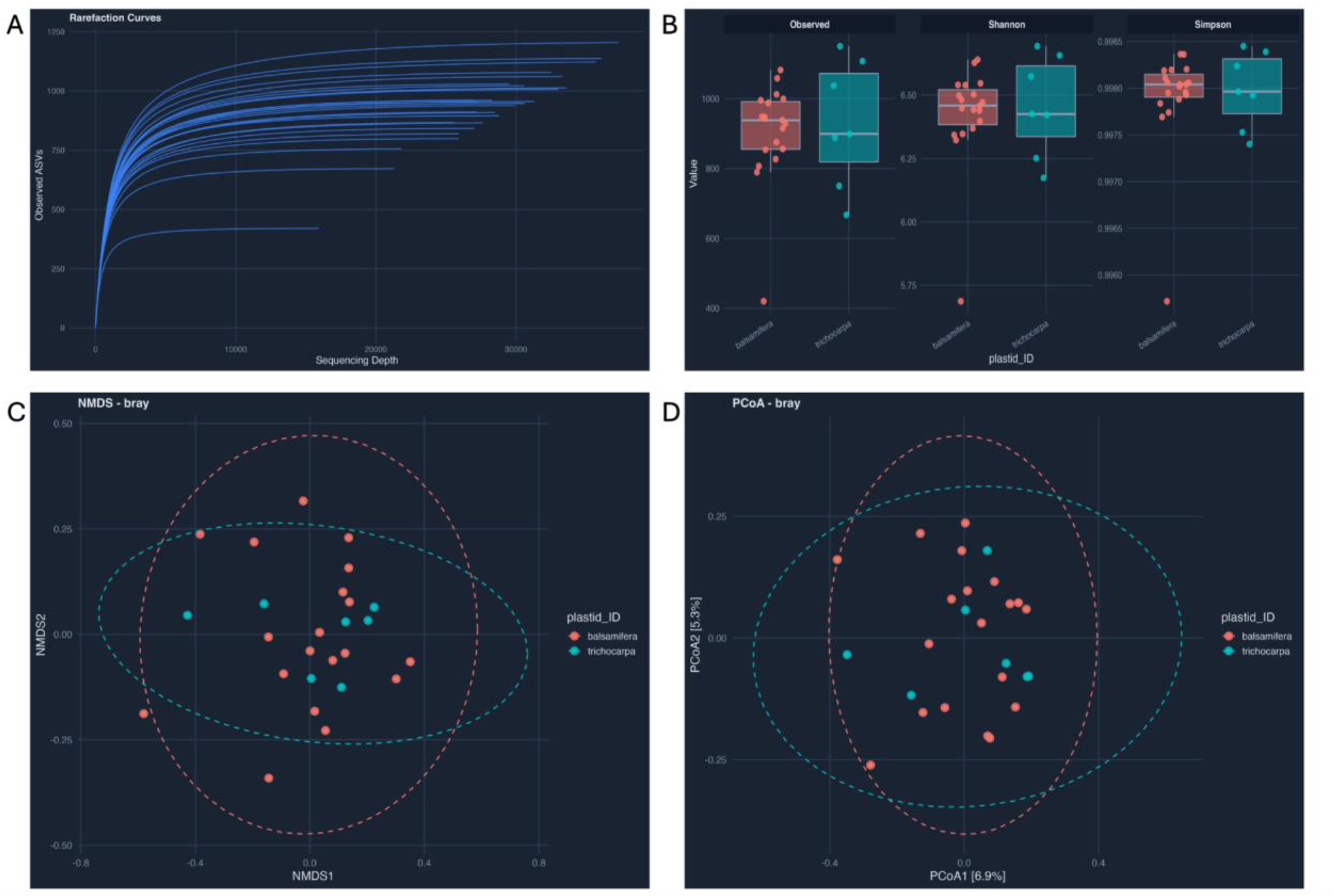
Downloaded plots of (A) Rarefaction (B) box plot of alpha diversity indices (Observed, Shannon, and Simpson), (C) NMDS ordination plot, and (D) PCoA ordination plot.

### 2.3 The ITS application (11 steps)

The ITS application extends the shared framework to an 11-step workflow that addresses the unique requirements of ITS amplicon sequencing. Unlike the 16S rRNA gene, the ITS region is highly variable in length (200-600 bp), which has two key consequences: fixed-length truncation is inappropriate because it would remove real ITS variants shorter than the truncation length, and primer removal is complicated by the possibility of read-through into the opposite primer when the amplified region is shorter than the read length.

The 11-step ITS workflow is: (1) Setup and file detection; (2) quality profile visualization of raw reads; (3) primer removal using cutadapt, including automatic N-prefiltering, primer orientation verification with a hit count table displayed for user confirmation, support for multiple forward and reverse primers, and automatic cutadapt detection or installation via pip; (4) quality filtering without truncation but with a configurable minimum length parameter (default 50 bp) to remove spuriously short sequences; (5-6) error learning, dereplication, merging, and chimera removal as in the 16S workflow; (7) taxonomy assignment using the UNITE database for fungal classification; and (8-11) phyloseq construction, visualization, PERMANOVA, and ANCOM-BC2, identical to the 16S application.

### 2.4 PERMANOVA analysis

Both applications include a dedicated PERMANOVA step using the vegan package (Dixon, 2003). Users enter a formula specifying one or more variables from their metadata (e.g., Treatment + Age), select a distance method (Bray-Curtis, Jaccard, or Euclidean), and configure the number of permutations (default 9,999). The application displays a live command preview showing the exact R commands that will be executed. Results include the adonis2 test table with all statistics rounded to three decimal places, a betadisper test for homogeneity of group dispersions with interpretive text, and automatic pairwise PERMANOVA with Bonferroni-adjusted p-values when more than two groups are present. All results are downloadable as CSV files.

### 2.5 ANCOM-BC2 differential abundance analysis

The final step in both applications implements ANCOM-BC2 for differential abundance analysis (Lin & Peddada, 2024). ANCOM-BC2 identifies taxa whose absolute abundances differ significantly between groups while correcting for both sample-specific and taxon-specific biases. The application uses the raw (untransformed) phyloseq object, as ANCOM-BC2 handles its own bias correction internally, and displays a notice informing users of this behavior.

Users specify fixed effects and optional random effects formulas as free-text input, select a grouping variable for global and pairwise tests, and configure the taxonomic level (default Genus), p-value adjustment method, prevalence filter, and library size cutoff. A collapsible advanced settings panel exposes all ANCOM-BC2 parameters including pseudo-count sensitivity analysis, structural zero detection, regularization, iteration and EM controls, mdFDR control, and optional Dunnett’s and trend tests. A live command preview shows the complete ancombc2() call. The analysis runs asynchronously with elapsed-time progress reporting.

Results include four outputs: (i) a primary analysis table showing which taxa are differentially abundant for each covariate in the model (ii) a global test table identifying taxa that are differentially abundant across any groups; (iii) a pairwise directional test table showing which taxa differ between each pair of groups with direction of change; and (iv) Dunnett’s type test results showing which taxa differ between each treatment group and a single control group. All tables are downloadable as CSV files.

### 2.6 Data transformation options

Both applications offer three data transformation methods applied after phyloseq construction: (i) rarefaction to even sequencing depth, defaulting to the minimum sample depth with user override, using phyloseq’s rarefy_even_depth function; (ii) relative abundance normalization, converting counts to proportions; and (iii) centered log-ratio (CLR) transformation via the microbiome package, which addresses compositionality and is recommended for use with Euclidean distance in ordination analyses (Aitchison distance). The transformation step includes a note recommending Euclidean distance when CLR is selected. All downstream visualizations and PERMANOVA use the transformed data, while ANCOM-BC2 always operates on the raw counts.

### 2.7 Session persistence and reproducibility

All pipeline state is automatically saved to the data directory after each step completes. On startup, the application checks for existing session files and offers to restore them with validation of file paths. Users navigating back to previously completed steps receive warnings that re-execution will invalidate downstream results. The complete R workspace can be exported as an .RData file and the phyloseq object as an .rds file, supporting integration with custom analysis workflows.

## 3. Conclusion and outlook

RAPID provides complete, code-free graphical interfaces for both 16S rRNA and ITS amplicon sequence analysis, bridging the gap between the powerful DADA2/phyloseq ecosystem and researchers without programming expertise. By offering two tailored applications that share a common architecture but address the specific requirements of each marker gene, RAPID eliminates the need for command-line interaction, script writing, or server infrastructure while preserving full access to the underlying algorithms.

Key architectural contributions include asynchronous processing that maintains interface responsiveness during long computations, automatic session persistence for multi-day analyses, guided step-by-step workflows that reduce parameter misconfiguration, and integration of statistical testing (PERMANOVA and ANCOM-BC2) within the same interface. All filtering and analysis parameters remain fully configurable, ensuring that experienced users retain control while novice users benefit from sensible defaults.

Future development directions include arbuscular mycorrhizal fungi analysis, agentic transformation of the apps, phylogenetic tree construction for UniFrac distance calculations, batch processing of multiple projects, integration of additional differential abundance methods, network analysis capabilities, and a unified launcher that allows users to select between the 16S and ITS workflows from a single entry point. Community contributions are welcome via the project’s open-source repository.

RAPID is designed to democratize access to rigorous amplicon sequence analysis for both bacterial and fungal communities, enabling a broader community of microbiome researchers to leverage the DADA2 algorithm’s strengths without compromising analytical rigor or reproducibility.

## References

Batut, B., Hiltemann, S., Bagnacani, A., Baker, D., Bhardwaj, V., Blank, C., Bretaudeau, A., Brillet-Guéguen, L., Čech, M., Chilton, J., Clements, D., Doppelt-Azeroual, O., Erxleben, A., Freeberg, M. A., Gladman, S., Hoogstrate, Y., Hotz, H.-R., Houwaart, T., Jagtap, P., … Grüning, B. (2018). Community-Driven Data Analysis Training for Biology. Cell Systems, 6(6), 752–758.e1. 10.1016/j.cels.2018.05.012

Bolyen, E., Rideout, J. R., Dillon, M. R., Bokulich, N. A., Abnet, C. C., Al-Ghalith, G. A., Alexander, H., Alm, E. J., Arumugam, M., Asnicar, F., Bai, Y., Bisanz, J. E., Bittinger, K., Brejnrod, A., Brislawn, C. J., Brown, C. T., Callahan, B. J., Caraballo-RodrÍguez, A. M., Chase, J., … Caporaso, J. G. (2019). Reproducible, interactive, scalable and extensible microbiome data science using QIIME 2. Nature Biotechnology, 37(8), 852–857. 10.1038/s41587-019-0209-9

Callahan, B. J., McMurdie, P. J., & Holmes, S. P. (2017). Exact sequence variants should replace operational taxonomic units in marker-gene data analysis. The ISME Journal, 11(12), 2639–2643. 10.1038/ismej.2017.119

Callahan, B. J., McMurdie, P. J., Rosen, M. J., Han, A. W., Johnson, A. J. A., & Holmes, S. P. (2016). DADA2: High-resolution sample inference from Illumina amplicon data. Nature Methods, 13(7), 581–583. 10.1038/nmeth.3869

Dixon, P. (2003). VEGAN, a package of R functions for community ecology. Journal of Vegetation Science, 14(6), 927–930. 10.1111/j.1654-1103.2003.tb02228.x

Hugerth, L. W., & Andersson, A. F. (2017). Analysing Microbial Community Composition through Amplicon Sequencing: From Sampling to Hypothesis Testing. Frontiers in Microbiology, 8. 10.3389/fmicb.2017.01561

Jia, L., Yao, W., Jiang, Y., Li, Y., Wang, Z., Li, H., Huang, F., Li, J., Chen, T., & Zhang, H. (2022). Development of interactive biological web applications with R/Shiny. Briefings in Bioinformatics, 23(1), bbab415. 10.1093/bib/bbab415

Lin, H., & Peddada, S. D. (2024). Multigroup analysis of compositions of microbiomes with covariate adjustments and repeated measures. Nature Methods, 21(1), 83–91. 10.1038/s41592-023-02092-7

Martin, M. (2011). Cutadapt removes adapter sequences from high-throughput sequencing reads. EMBnet.Journal, 17(1), 10–12. 10.14806/ej.17.1.200

McMurdie, P. J., & Holmes, S. (2013). phyloseq: An R Package for Reproducible Interactive Analysis and Graphics of Microbiome Census Data. PLOS ONE, 8(4), e61217. 10.1371/journal.pone.0061217

McMurdie, P. J., & Holmes, S. (2015). Shiny-phyloseq: Web application for interactive microbiome analysis with provenance tracking. Bioinformatics, 31(2), 282–283. 10.1093/bioinformatics/btu616

Murali, A., Bhargava, A., & Wright, E. S. (2018). IDTAXA: A novel approach for accurate taxonomic classification of microbiome sequences. Microbiome, 6, 140. 10.1186/s40168-018-0521-5

Nilsson, R. H., Larsson, K.-H., Taylor, A. F. S., Bengtsson-Palme, J., Jeppesen, T. S., Schigel, D., Kennedy, P., Picard, K., Glöckner, F. O., Tedersoo, L., Saar, I., Kõljalg, U., & Abarenkov, K. (2019). The UNITE database for molecular identification of fungi: Handling dark taxa and parallel taxonomic classifications. Nucleic Acids Research, 47(D1), D259–D264. 10.1093/nar/gky1022

Quast, C., Pruesse, E., Yilmaz, P., Gerken, J., Schweer, T., Yarza, P., Peplies, J., & Glöckner, F. O. (2013). The SILVA ribosomal RNA gene database project: Improved data processing and web-based tools. Nucleic Acids Research, 41(Database issue), D590–596. 10.1093/nar/gks1219

Schoch, C. L., Seifert, K. A., Huhndorf, S., Robert, V., Spouge, J. L., Levesque, C. A., Chen, W., Fungal Barcoding Consortium, Fungal Barcoding Consortium Author List, Bolchacova, E., Voigt, K., Crous, P. W., Miller, A. N., Wingfield, M. J., Aime, M. C., An, K.-D., Bai, F.-Y., Barreto, R. W., Begerow, D., … Schindel, D. (2012). Nuclear ribosomal internal transcribed spacer (ITS) region as a universal DNA barcode marker for Fungi. Proceedings of the National Academy of Sciences, 109(16), 6241–6246. 10.1073/pnas.1117018109

Wang, Q., Garrity, G. M., Tiedje, J. M., & Cole, J. R. (2007). Naive Bayesian classifier for rapid assignment of rRNA sequences into the new bacterial taxonomy. Applied and Environmental Microbiology, 73(16), 5261–5267. 10.1128/AEM.00062-07

Zheng, J.-J., Wang, P.-W., Huang, T.-W., Yang, Y.-J., Chiu, H.-S., Sumazin, P., & Chen, T.-W. (2022). MOCHI: A comprehensive cross-platform tool for amplicon-based microbiota analysis. Bioinformatics, 38(18), 4286–4292. 10.1093/bioinformatics/btac494

Zhou, J., He, Z., Yang, Y., Deng, Y., Tringe, S. G., & Alvarez-Cohen, L. (2015). High-Throughput Metagenomic Technologies for Complex Microbial Community Analysis: Open and Closed Formats. mBio, 6(1), 10.1128/mbio.02288-14. https://doi.org/10.1128/mbio.02288-14

