## Supplementary Figures for "RAPID: an interactive R/Shiny platform for end-to-end 16S rRNA and ITS amplicon sequence analysis using DADA2"

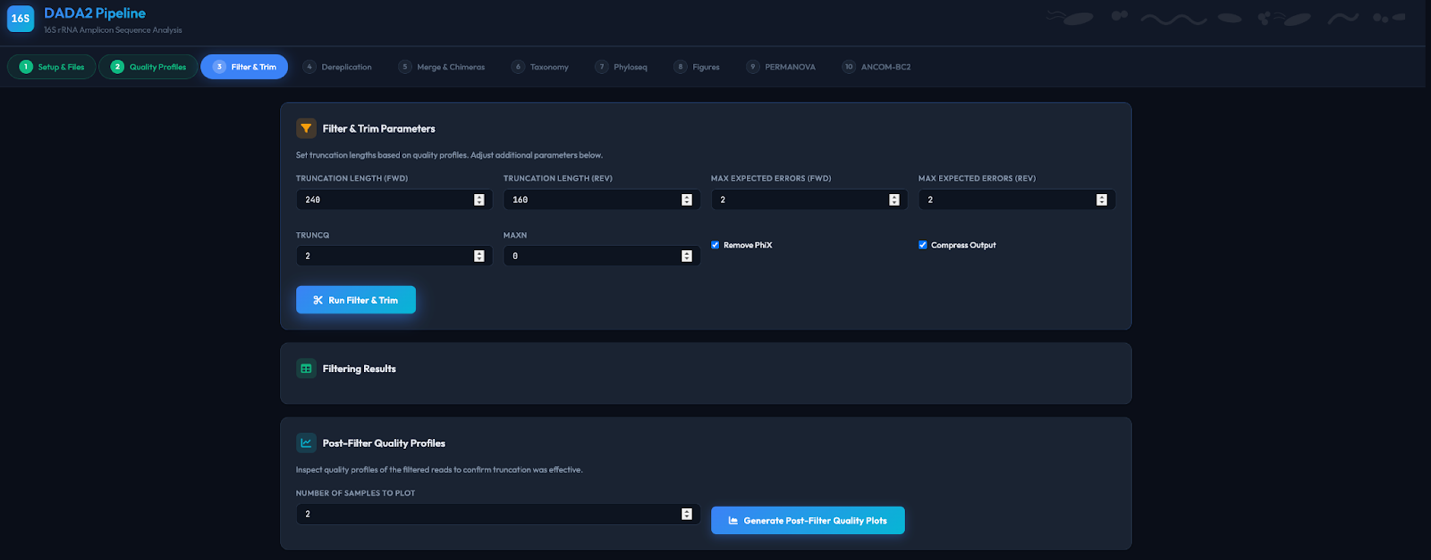

**Supplementary Figure 1.** Screenshot of the filter and trim step with configurable parameters and sensible defaults based on standard DADA2 16S workflow.

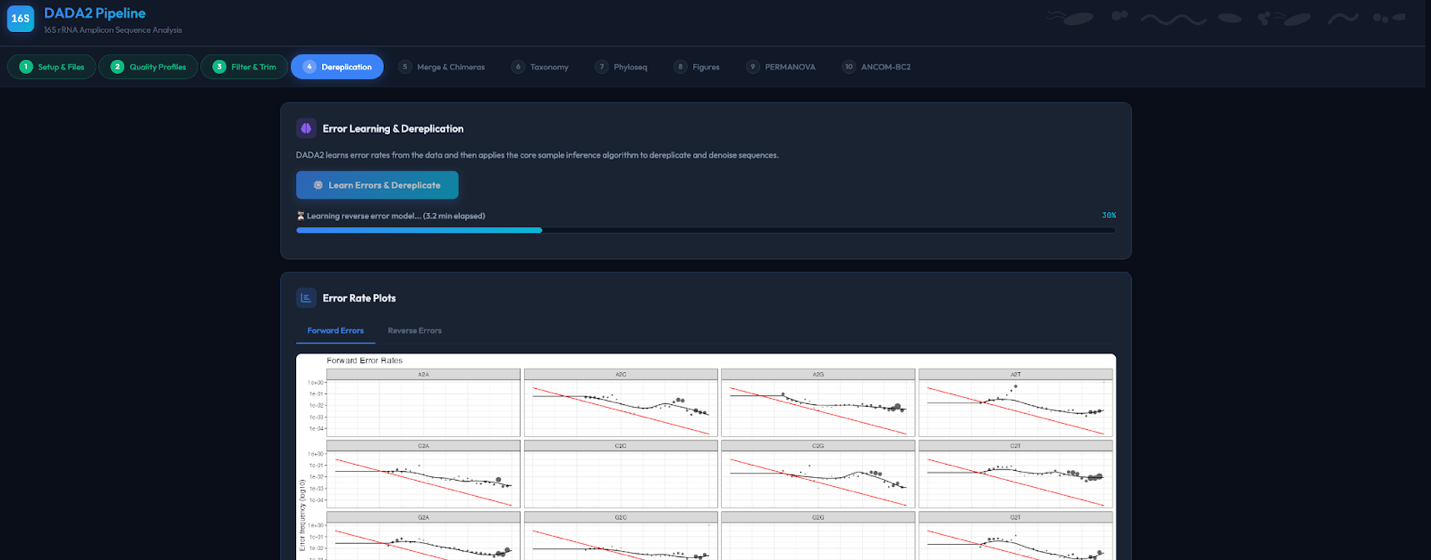

**Supplementary Figure 2.** Screenshot of the DADA2 error learning algorithm and dereplication step.

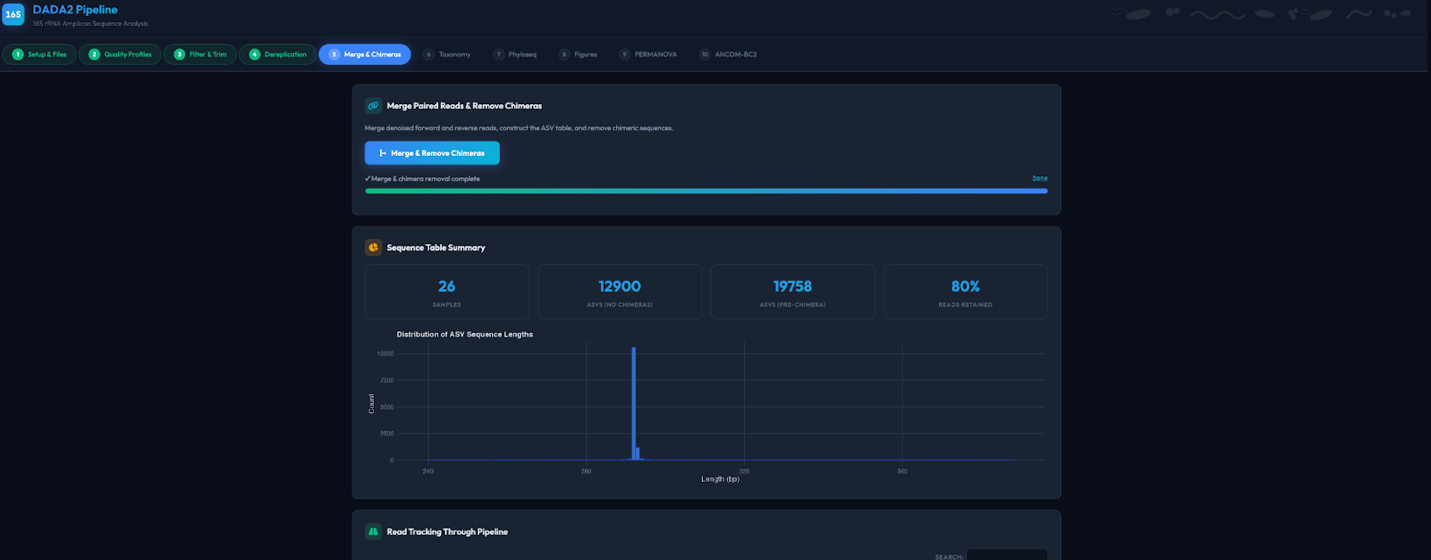

**Supplementary Figure 3.** Screenshot of the merged read pairs and remove chimera step.

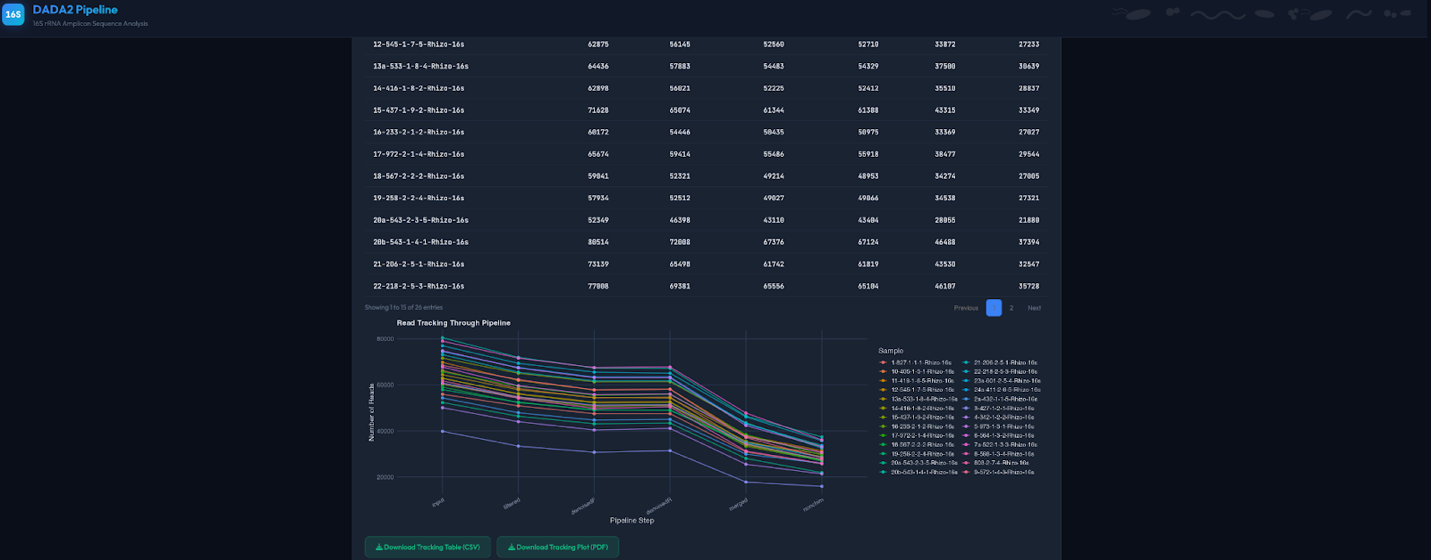

**Supplementary Figure 4.** Screenshot of the read tracking table and line graph for each sample.

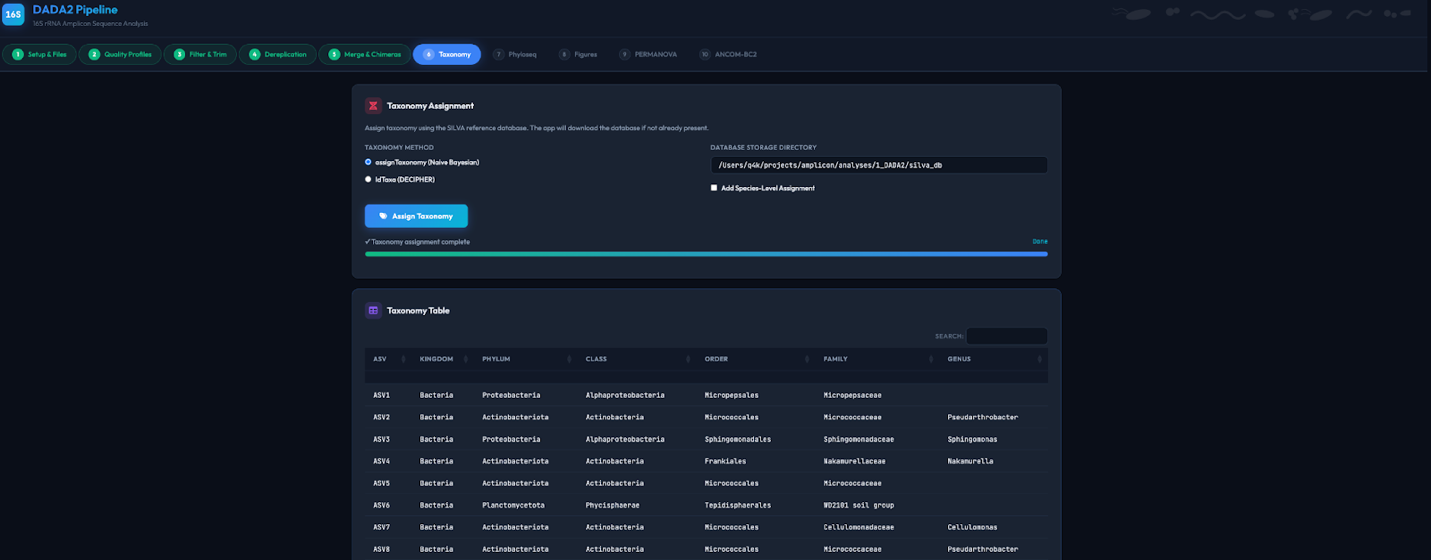

**Supplementary Figure 5.** Screenshot of the assignTaxonomy step using Bayesian or IDTAXA classifier algorithm.

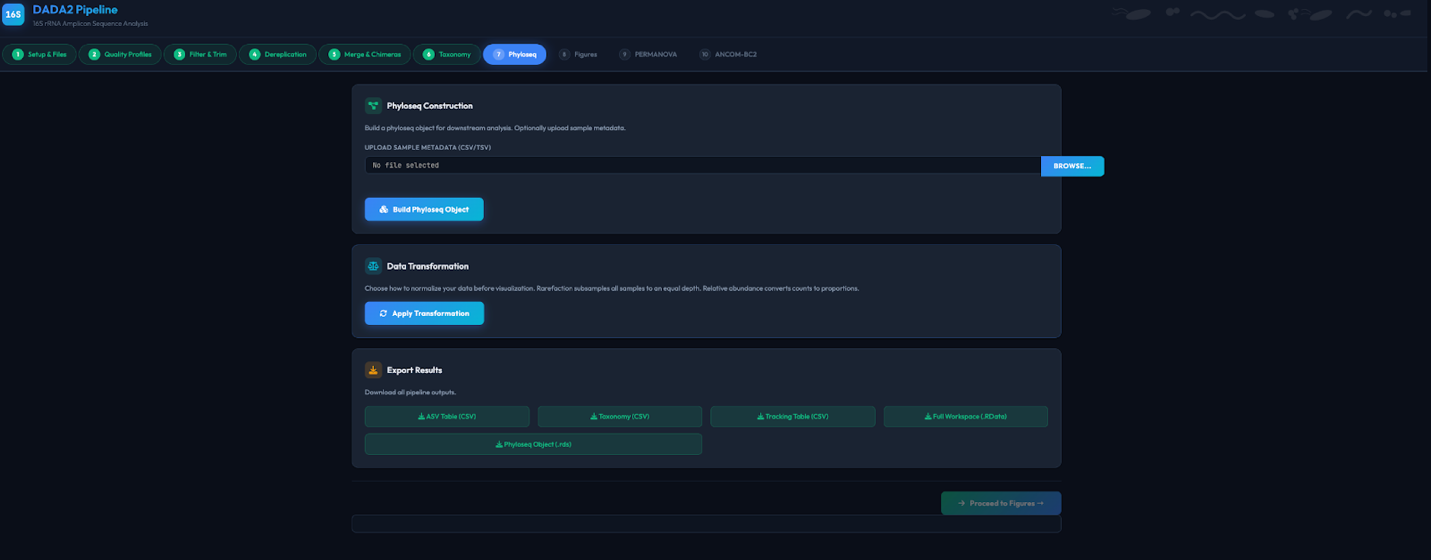

**Supplementary Figure 6.** Screenshot of the Phyloseq step where users upload metadata file for phyloseq object construction.

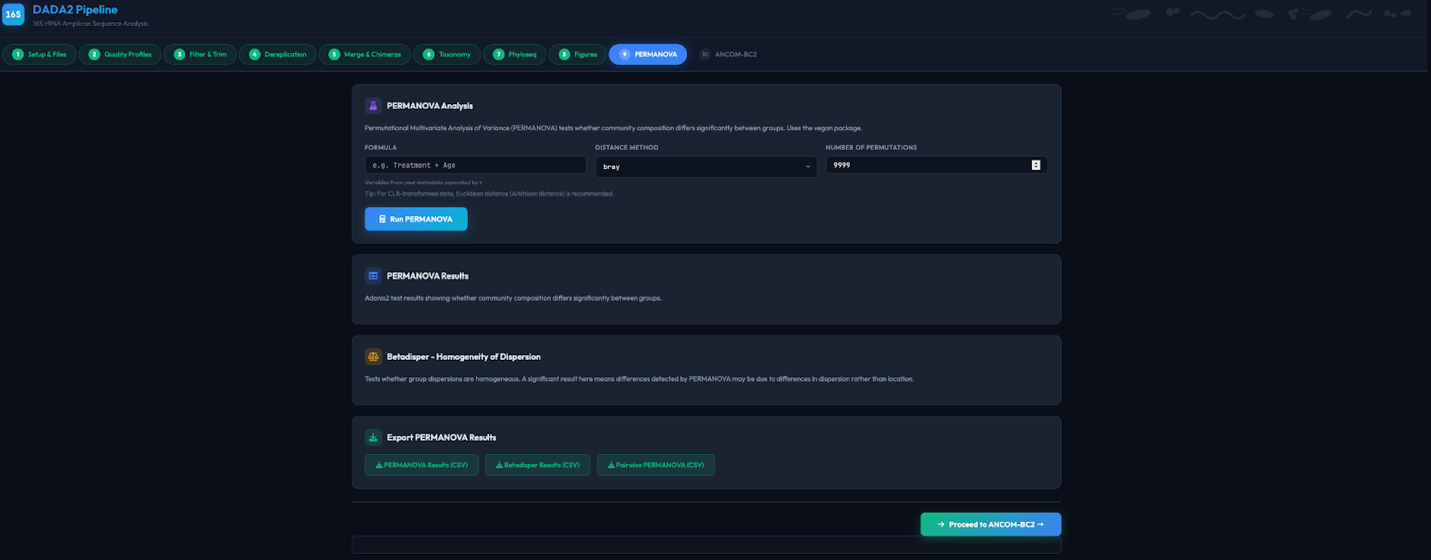

**Supplementary Figure 7.** Screenshot of the PERMONOVA step implemented using the vegan package.

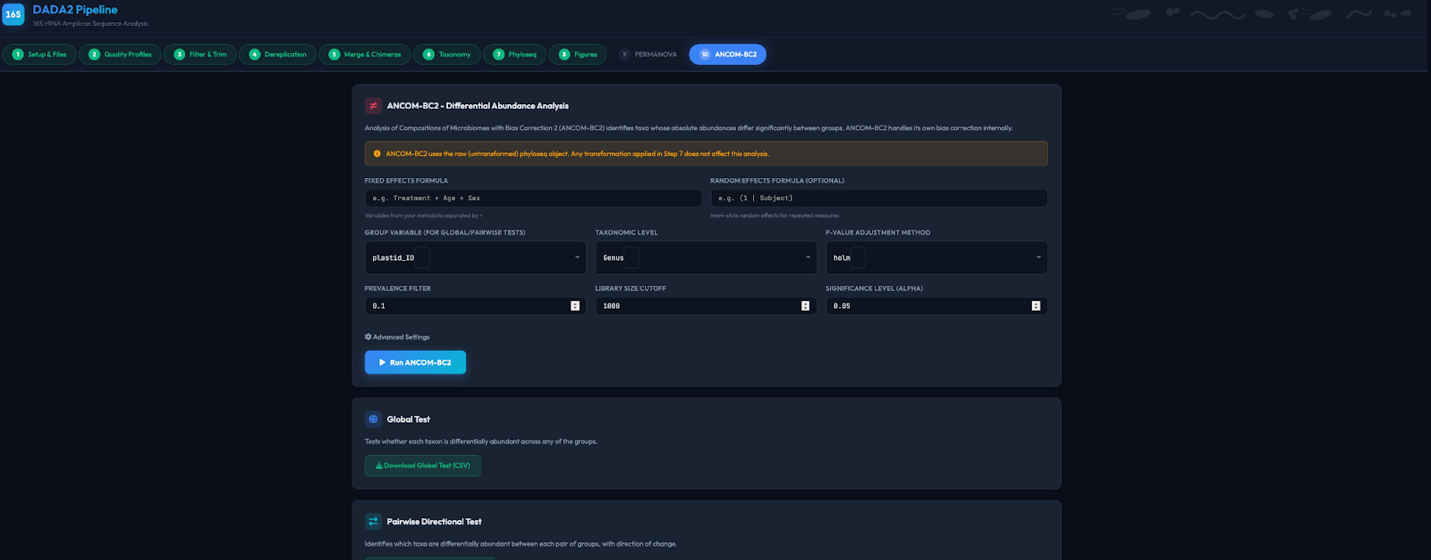

**Supplementary Figure 8.** Screenshot of the ANCOMBC-2 step for the differential abundance analysis at every taxa.
